# Hsp27 reduces glycation-induced toxicity and aggregation of α-synuclein

**DOI:** 10.1101/2020.03.03.975037

**Authors:** Hugo Vicente Miranda, Ana Chegão, Márcia Oliveira, Bárbara Fernandes Gomes, Francisco J. Enguita, Tiago Fleming Outeiro

## Abstract

α-synuclein (aSyn) is a major player in Parkinson’s disease (PD) and a group of other disorders collectively known as synucleinopathies, but the precise molecular mechanisms involved are still unclear. aSyn, as virtually all proteins, undergoes a series of posttranslational modifications (PTMs) during its lifetime which can affect its biology and pathobiology. We recently showed that glycation of aSyn by methylglyoxal (MGO) potentiates its oligomerization and toxicity, induces dopaminergic neuronal cell loss in mice, and affects motor performance in flies. Small heat-shock proteins (sHsps) are molecular chaperones that facilitate the folding of proteins or target misfolded proteins for clearance. Importantly, sHsps were shown to prevent aSyn aggregation and cytotoxicity. Upon treating cells with increasing amounts of methylglyoxal, we found that the levels of Hsp27 decreased in a dose-dependent manner. Therefore, we hypothesized that restoring the levels of Hsp27 in glycating environments could alleviate the pathogenicity of aSyn. Consistently, we found that Hsp27 reduced MGO-induced aSyn aggregation in cells, leading to the formation of non-toxic aSyn species. Remarkably, increasing the levels of Hsp27 suppressed the deleterious effects induced by MGO. Our findings suggest that in glycating environments, the levels of Hsp27 are important for modulating the glycation-associated cellular pathologies in synucleinopathies.

## Introduction

α-synuclein (aSyn) is a small protein of 140 amino acid residues abundant in the brain and in various other tissues. In the brain, it is abundant in the presynaptic compartment, but exists also in the nucleus of neurons ^1–5^. aSyn is the major component of pathognomonic protein inclusions, known as Lewy bodies and Lewy neurites, present in the brains of patients affected by Parkinson’s disease (PD) or dementia with Lewy bodies ^6–8^. It is also present in glial cytoplasmic inclusions in multiple system atrophy (MSA)^6–8^. Together, diseases associated with the aggregation of aSyn in the brain are known as synucleinopathies ^6–10^. However, the molecular mechanisms underlying aSyn aggregation are still elusive.

aSyn is an intrinsically disordered protein and is prone to aggregation under various conditions ^11,12^. The process of aSyn aggregation has been the subject of intensive study, and is thought to proceed through the assembly of the protein into dimers, oligomers, and fibrils, which may then accumulate in protein inclusions ^9,12–14^.aSyn undergoes several posttranslational modifications (PTMs) that can modulate its aggregation and cytotoxicity ^15^. aSyn is a long-lived protein that while it does not contain any arginine residues, it is lysine-rich, making it a preferably target for glycation in the lysine residues ^16,17^. Several studies suggest that glycation might be an important contributor to the pathobiology of aSyn ^17,18^. In a previous study, we found that aSyn is glycated in the brains of PD patients and that this modification modulates aSyn pathogenicity, contributing to the neurodegenerative process ^19^. Furthermore, methylglyoxal (MGO), the strongest glycating agent in living cells, exacerbates aSyn aggregation and toxicity in several models of synucleinopathies, such as yeast, human cells, and animal models of PD ^19^. In addition to inducing aSyn oligomerization, MGO also affects the correct clearance of aSyn by impairing the activities of the proteasome and autophagy clearance systems ^19^.

The small heat-shock proteins (sHsps) are a family of small molecular weight proteins that are characterized by a highly conserved domain, known as alpha-crystallin domain ^20,21^. sHsps play an important role in cellular proteostasis ^22–25^, in particular by preventing protein aggregation. They act as molecular chaperones, promoting protein folding/refolding, stabilization, and by targeting damaged proteins for degradation ^24–26^. Furthermore, sHsps are stress response proteins, upregulated under unfavorable conditions such as heat-shock, metabolic, oxidative, and chemical stress ^27,28^. However, sHsps are not only involved in stress responses, but also in stress tolerance, cell death, differentiation, cell cycle, and signal transduction ^29^. sHsps have also been reported to play a role in aging and protein aggregation disorders, such as neurodegenerative diseases like Alzheimer’s disease and PD ^30–34^. Remarkably, Hsp27, one of the members of the sHsp family, is present in Lewy bodies ^35,36^.

Hsp27 is ubiquitously expressed, at highest levels in the skeletal, smooth, and cardiac muscles ^37^. Like other sHsps, it can assemble in smaller oligomers, such as dimers or tetramers, or in large oligomeric complexes ^38–42^. Hsp27 is a target of PTMs, such as phosphorylation, which induces the shift in the oligomeric status of Hsp27 ^40^. Hsp27 can have different functions, depending on its oligomerization state: in large oligomers, it acts mainly as a molecular chaperone, also exhibiting anti-apoptotic properties ^42,43^; in small oligomers, Hsp27 has reduced chaperone activity and acts in the dynamics of microfilaments, contributing to the stabilization of F-actin filaments ^44,45^. The role of Hsp27 in protein aggregation is well documented. The levels of brain Hsp27 brain are increased in the PD brain (cortex) ^46,47^, and is also present in Lewy bodies and co-localizes with aSyn ^35,36^. In vitro, Hsp27 binds to aSyn preventing aggregation and fibril elongation, and also reducing aSyn cytotoxicity in a concentration-dependent manner ^23,35,48–50^.

Like aSyn, Hsp27 can also be glycated by MGO ^51–55^. Interestingly, this modification promotes the formation of large oligomeric complexes of Hsp27, enhancing its chaperone activity and anti-apoptotic properties ^51,55^. Moreover, high levels of glucose increase Hsp27 glycation and reduce its levels in a dose dependent manner ^56^.

Here, we report that MGO reduces the levels of Hsp27 in a dose dependent manner. Therefore, we hypothesized that restoring the levels of Hsp27 could prevent glycation-associated aSyn pathobiology. Consistently, Hsp27 overexpression reduced the deleterious effects of MGO in a cellular model of synucleinopathies. Our findings suggest that Hsp27 might constitute an important therapeutic target for modulating glycation-induced pathogenesis of aSyn.

## Materials and Methods

### Cell culture

Human H4 neuroglioma cells were maintained at 37°C in OPTI-MEM I (Gibco, Invitrogen, CA, USA) supplemented with 10% fetal bovine serum (FBS) (Gibco, Invitrogen, CA, USA). H4 neuroglioma cells were seeded in 35 mm imaging dishes (Ibidi, 170.000 cells/ dish) for immunocytochemistry assays, 12-well plate (TPP, 90.000 cells/ well) for cytotoxicity assays, 6-well plate (TPP, 190.000 cells/ well) for protein analysis, or 6 cm plates (TPP, 300.000 cells/ dish) for triton-X 100 solubility assay.

### Analysis of protein glycation and Hsp27 profile

H4 neuroglioma naïve cells were plated and treated 24 hours later with increasing amounts of MGO (0.2; 0.5; 0.75; 1 mM), prepared as previously ^19^. 24 hours after treatment, cells were collected and protein extracts prepared and quantified as previously ^19,57^. Protein glycation profile was assessed by western blotting, probing for argpyrimidine (a kind gift from K. Uchida, Laboratory of Food and Biodynamics, Nagoya University Graduate School of Bioagricultural Sciences, Japan) as previously ^58^. Hsp27 levels were probed with anti-Hsp27 (F-4 1:2000; Santa Cruz Biotechnology, Dallas, TX, USA). Detection procedures were performed according to ECL system (GE Healthcare, Life Sciences; Little Chalfont, UK), and the signal was detected using a ChemiDoc^™^ Imaging Systems (Bio-Rad, Hercules, CA, USA). Densitometry was performed using ImageJ - Image Processing and Analysis in Java. When required, membranes were incubated with stripping solution (250 mM Glycine, 0.1 % of 10 % SDS, pH 2.0) for 45 minutes at room temperature with agitation, followed by 4 washing steps, twice with 1x TBS and twice with 1x TBS supplemented with 10% Tween 20 solution. Membranes were then incubated with blocking solution for 30 minutes, before reprobing.

### Mass spectrometry

H4 neuroglioma cells protein extracts were separated by SDS-PAGE and the correspondent 25 kDa fragment was excised from the gel. Peptide mass fingerprinting analysis was performed as previously ^19,57^, in an Applied Biosystems 4700 Proteomics Analyzer with TOF/TOF ion optics. A double miscleavage was allowed, and oxidation of methionyl residues, acetylation of the N-terminal region, and carboxyethylation (CEL) of lysine residues, as well as argpyrimidine or hidroimidazolones formation at arginine residues were assumed as variable modifications. All peaks with S/N 4-5 were included, with a taxonomic restriction to *Homo sapiens*. The criteria used to accept the identification were significant homology scores achieved in Mascot (p<0.05).

### Expression and purification of recombinant aSyn and Hsp27

Human aSyn was expressed and purified as we previously described ^19^. SDS-PAGE, followed by western blotting analysis (using standard procedures), confirmed the monomeric purification of aSyn (anti-a-synuclein dilution 1:1000; BD Transduction Laboratories™, San Jose, CA, USA).

For human Hsp27, the *E. coli* strain BL-21 was transformed by heat shock with HSP27 PET16b construct (a kind gift from Dr. Paul Muchowski) and expression induced for 3 hours with IPTG (0.3 mM). Cells were pelleted and re-suspended in lysis buffer, supplemented with 10 mg/mL of lysozyme (Sigma-Aldrich, MO, USA). The cell suspension was then incubated on ice with constant stirring for 20 minutes, followed by the addition of 0.33 mL/L benzonase (Sigma-Aldrich, MO, USA), and subsequently incubated at room temperature for 20 minutes with constant stirring. Insoluble cellular debris were removed by centrifugation. 53 mL/L of a 200 mM dithiothreitol (Sigma-Aldrich, MO, USA) solution was added to the soluble supernatant, followed by incubation at room temperature with constant stirring for 10 minutes. Insoluble contaminants were removed by centrifugation and the supernatant was filtered with a 0.22 μm filter. The sample was loaded into an ion-exchange chromatography Q Sepharose TM fast flow column equilibrated with 20 mM Tris-HCl, pH 8.0. Proteins were eluted with a linear NaCl gradient (0 - 1.0 M), at a flow rate of 1.5 mL/min and the elution monitored at 280 nm. Protein-containing fractions were collected and probed by SDS-PAGE analysis using Coomassie staining. Fractions containing the Hsp27 were collected, concentrated by centrifugation using Amicon filters and applied to a gel filtration Superdex 75 column, equilibrated with 20 mM Tris-HCl buffer, pH 7.4, containing 100 mM NaCl. Proteins were eluted with the same buffer at a flow rate of 1 mL/min. Fractions containing Hsp27 were collected and concentrated by centrifugation using Amicon filters. SDS-PAGE, followed by western blotting analysis, confirmed the monomeric purification of Hsp27 (anti-Hsp27 1:2000; Santa Cruz Biotechnology, Dallas, TX, USA).

### In vitro aSyn aggregation

Recombinant monomeric aSyn was diluted at 140 μM in 30 mM Tris-HCl pH 7.4. Aggregation was induced as previously described ^19^. aSyn aggregation was evaluated alone or in the presence of Hsp27 (0.45 or 3 μM), and or in the presence of MGO (0.5 mM) or corresponding vehicle.

### Thioflavin T binding assay

The formation of amyloid fibrils was monitored by Thioflavin T (ThT) binding assay as previously described ^19^. Emission wavelength scan at 490 nm was performed with an excitation wavelength of 450 nm using a plate reader (Tecan Infinite 200, Männedorf, Switzerland).

### Native-PAGE electrophoresis

15 μg of total protein extract from purified proteins were separated by native electrophoresis using a Tetra cell (Bio-Rad; Hercules, CA, USA), in 12% polyacrylamide separation gel and a 4% polyacrylamide stacking gel in non-denaturing conditions, applying a constant voltage of 120 V.

### aSyn cytotoxicity assays

To assess exogenous aSyn toxicity, cells were plated and treated 24 hours later with recombinant aSyn species, either glycated or not, alone or in the presence of Hsp27 (0.45 or 3 μM). Cells were collected 24 hours later, and cytotoxicity was determined by LDH cytotoxicity assay (see below).

### Hsp27 overexpression

24h post-seeding, H4 neuroglioma cells were transfected with pcDNA3.1 vectors containing aSyn WT ^36^, SynT ^36^, or Hsp27 ^35^ alone, or co-transfected with aSyn WT and Hsp27, or SynT and Hsp27 pcDNA3.1 vectors, using FuGENE^®^ 6 (Roche, Basel, Switzerland), using standard procedures. 20 hours posttransfection, cells were treated with MGO (0.2 mM). 24 hours later, a second MGO treatment (0.2 mM) was performed. Cells were collected for analysis 48 hours post-transfection.

### aSyn, Hsp27 and glycation levels

H4 protein extracts were analyzed by western-blotting probing for aSyn (anti-a-synuclein dilution 1:1000; BD Transduction Laboratories™, San Jose, CA, USA), Hsp27 (anti-Hsp27 F-4 1:2000; Santa Cruz Biotechnology, Dallas, TX, USA), anti-MGO-derived Advanced Glycation and Products (AGEs) (KAL-KH001 Cosmo-Bio, USA, 1:500 dilution) and normalized to ²-actin levels (anti-b-actin 1:5000; Thermo Fisher Scientific; Waltham, MA, USA).

### LDH cytotoxicity assay

Cytotoxicity was measured using lactate dehydrogenase (LDH) kit (Clontech; Mountain View, CA, USA), according to manufacturer’s instructions.

### Triton-X 100 solubility assay

The solubility assays were performed as previously described ^19^.

### Immunocytochemistry

Immunocytochemistry was performed as previously described ^19^. Microscopy images were acquired in a Widefield fluorescent microscope Zeiss Axiovert 40 CFL (Carl Zeiss MicroImaging).

## Results

### MGO induces the formation of argpyrimidine modifications and reduces the levels of Hsp27

First, we determined the profile of argpyrimidine-modified proteins in human H4 cells induced by increasing concentrations of MGO (0.2; 0.5; 0.75 mM) after 24 hours of treatment. Argpyrimidine is a specific MGO-derived AGE at arginine residues. We identified a protein migrating with an apparent molecular weight of ~25 kDa **(Fig. 1 *A*)**. Surprisingly, that signal decreased in an MGO concentration-dependent manner, in contrast to the effect on other proteins **(Fig. 1 *A*)**. To identify this protein, cell extracts were separated by SDS-PAGE and the correspondent ~25 kDa fragment was excised from the gel and processed for mass spectrometry analysis, resulting in the identification of Hsp27 (55% sequence coverage).

**Figure 1.**
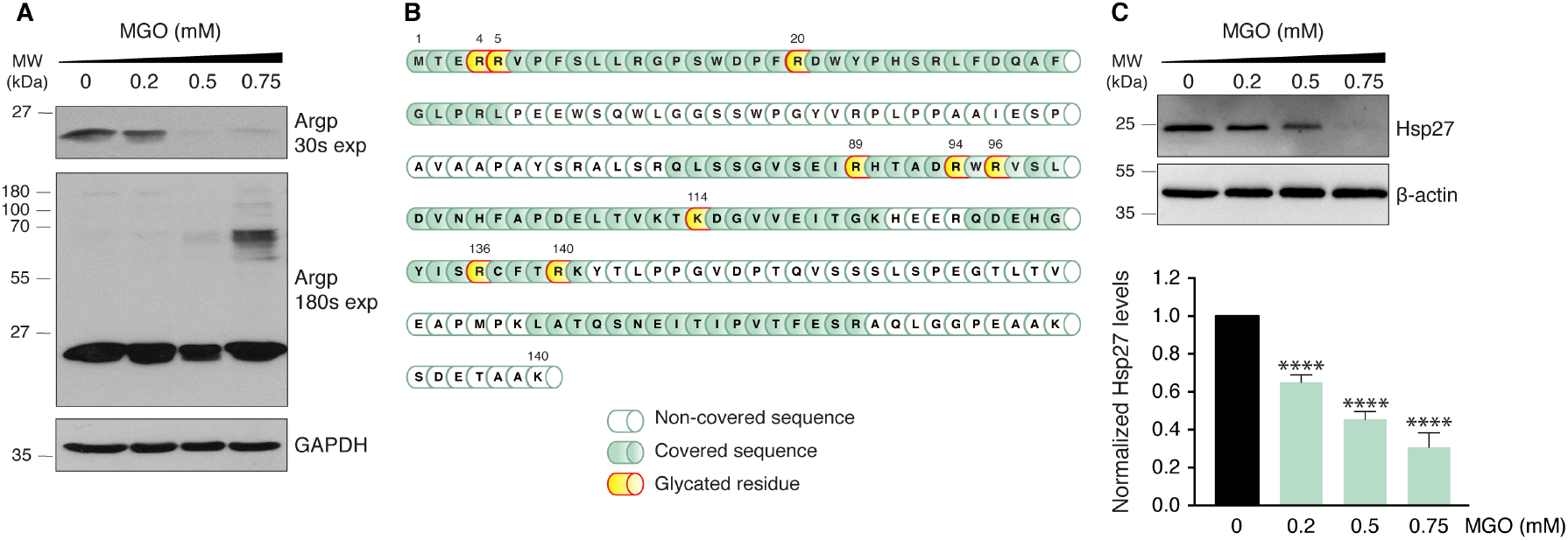
The levels of Hsp27 decrease in an MGO-dependent manner. H4 cells were treated with increasing concentrations of MGO for 24h. **(*A*)** Protein extracts were separated by SDS-PAGE and were immunoblotted with anti-Argpyrimidine and anti-GAPDH antibodies (loading control). Two exposure times are shown (30 or 180 seconds of signal acquisition in ChemiDoc^™^ Touch Imaging System, Bio-Rad). **(*B*)** A gel fragment at ~25 kDa was cut and processed for mass spectrometry analysis, leading to the indentification of Hsp27. Hsp27 schematics show the different glycated residues (yellow), with a 55% sequence coverage (dark green). **(*C*)** Protein extracts were probed for Hsp27 and β-actin (loading control). Data presented as fold ratio to vehicle-treated cells (n=3). **** *p* < 0.0001, oneway ANOVA, followed by Tukey’s multiple comparisons test.

We identified several glycated residues in Hsp27, mainly in arginine residues (**Fig. 1 *B*** and **Supplemental table 1**), and immunoblot analysis confirmed that the ~25 kDa protein was recognized by an antibody specific for Hsp27. Interestingly, we confirmed that the levels of Hsp27 decreased in cells treated with increasing concentrations of MGO (**Fig. 1 *C***).

### Hsp27 reduces aSyn aggregation and cytotoxicity

Hsp27 was previously shown to reduce the oligomerization of aSyn ^59^. However, the effect of Hsp27 on the oligomerization of glycated aSyn has not been investigated. To address this, we tested the effect of sub-stoichiometric levels of Hsp27 (300:1 or 300:7, aSyn:Hsp27) on aSyn aggregation (140 μM), in the presence or absence of glycation conditions. To study the kinetics of aggregation, aSyn was incubated under controlled temperature and shaking conditions, and samples were collected at specific timepoints. Using native-PAGE electrophoresis followed by immunoblot analysis, we observed that, with time, aSyn species appeared at the top of the gel (**Fig. 2 *A***), indicating the formation of high molecular weight (HMW) species ^60–64^. In the presence of Hsp27, the formation of these HMW species was reduced (**Fig. 2 *A***).

As we and others previously reported that while suppressing fibril formation, glycation of aSyn by MGO increases its oligomerization ^19,65^. Here, we evaluated if Hsp27 protects from glycation effects on aSyn oligomerization and found that it reduced the formation of HMW species of aSyn (**Fig. 2 *B***).

**Figure 2.**
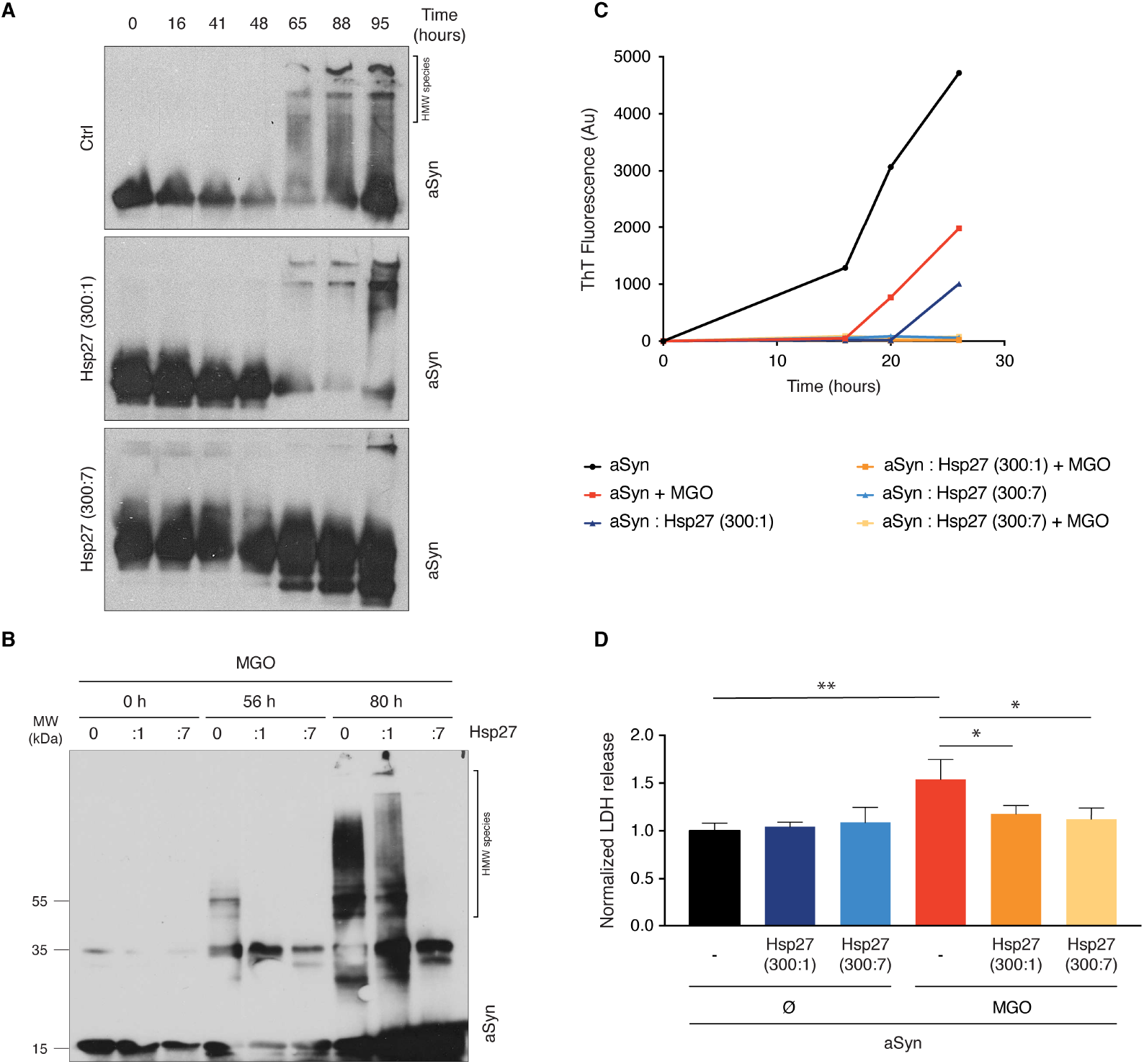
Hsp27 decreases aSyn aggregation and toxicity in a dose dependent manner. aSyn aggregation kinetics (140 μM) in the absence or presence of Hsp27 (300:1 or 300:7, aSyn:Hsp27) was evaluated by **(*A*)** native-PAGE followed by immunoblotting with an anti-aSyn antibody. **(*B*)** SDS-PAGE followed by immunoblotting with an anti-aSyn antibody, in control or MGO-glycation conditions. **(*C*)** ThT aggregation assays. **(*D*)** The cytotoxicity of the resulting species was evaluated in H4 cells using LDH release as a readout. Data normalized to control conditions without Hsp27 or MGO (n=3). * *p* < 0.05, ** *p* < 0.01, one-way ANOVA, followed by Tukey’s multiple comparisons test.

In particular, at a 300:7 (aSyn:Hsp27) molar ratio, aSyn remains mainly monomeric or as lower molecular weight species, suppressing the formation of HMW species, even at 80h of incubation. We also confirmed a reduction in the formation of aSyn amyloid structures in the presence of Hsp27, using Thioflavin T (ThT) as a readout (**Fig. 2 *C***).

Next, to determine whether Hsp27 affected the toxicity of glycated aSyn species, we exposed H4 cells to those species, for 24h, and assessed the overall membrane integrity (LDH assay) as a readout of cytotoxicity. While aSyn exposed to MGO was toxic, as we have previously shown, the species incubated with Hsp27 were not (**Fig. 2 *D***).

### Hsp27 reduces MGO-associated aSyn pathobiology in cellular models

Next, we investigated the effects of Hsp27 on MGO-associated aSyn cellular pathologies in cells. Briefly, cells expressing aSyn alone or together with Hsp27 were treated with MGO 20 hours post-transfection, and then received a second treatment 24 hours later, in order to ensure glycation conditions were maintained. Protein extracts were then prepared and analyzed by immunoblot analyses. In contrast to the effect of MGO observed on the endogenous levels of Hsp27 (**Fig. 1**), we found that in cells overexpressing Hsp27 the levels did not decrease (**Fig. 3 *A-D***), while the general glycation increased in an MGO-dependent manner (**Fig. 3 *EH***). In cells expressing aSyn WT, we observed that Hsp27 prevented aSyn cytotoxicity by ~10%. Impressively, Hsp27 overexpression also prevented the MGO-associated increase of aSyn levels (~ 56%) (**Fig. 4 *A***) and cytotoxicity (~ 28%) (**Fig. 4 *B***).

**Figure 3.**
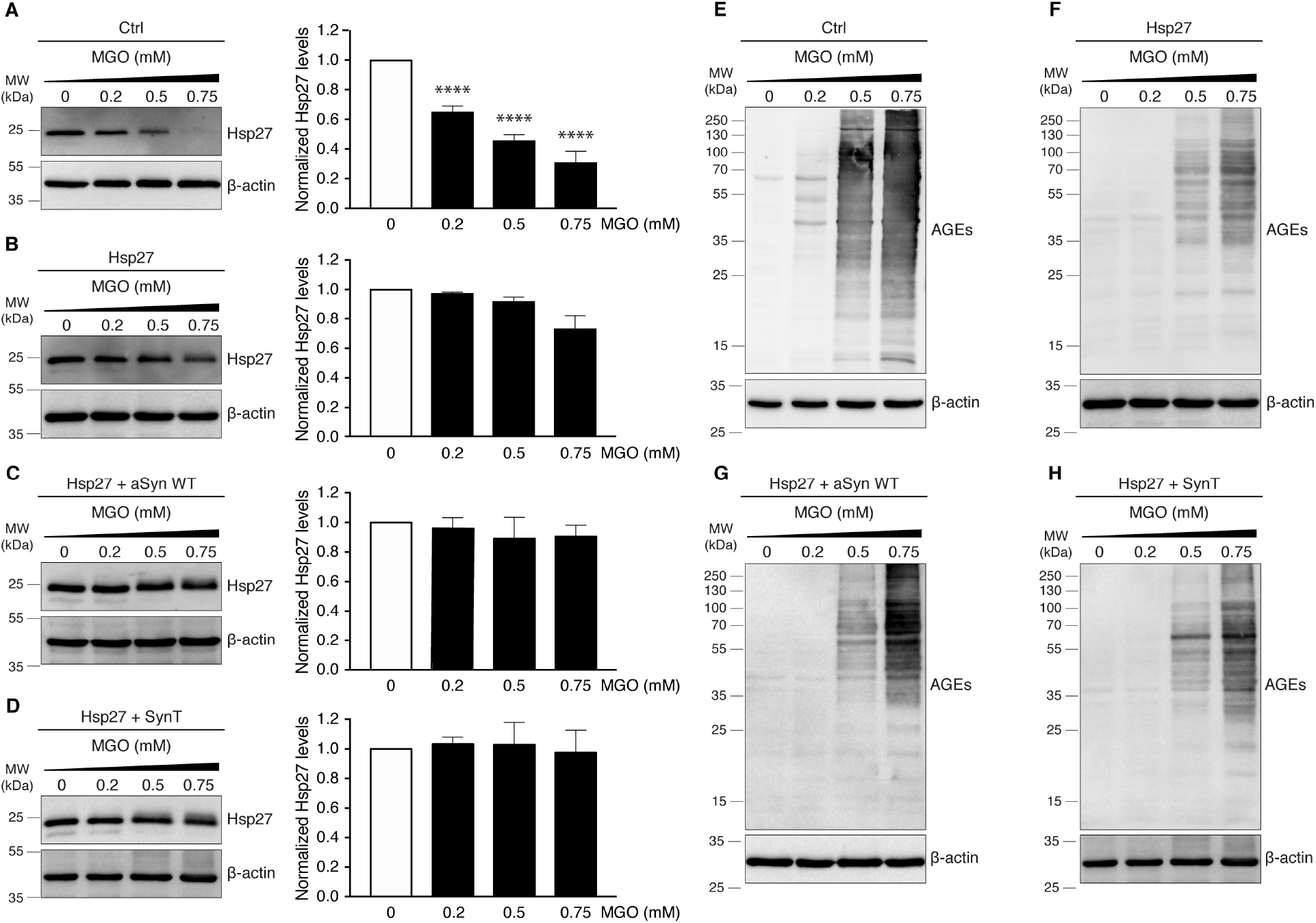
Hsp27 overexpression compensates for MGO-induced decrease of Hsp27 levels. H4 cells were treated with increasing concentrations of MGO for 24h. Protein extracts from **(*A, E*)** naïve cells; or from cells overexpressing **(*B, F*)** Hsp27; **(*C, G*)** Hsp27 and aSyn; or **(*D, H*)** Hsp27 and SynT, were separated by SDS-PAGE and immunoblotted for Hsp27 **(*A-D*)** or AGEs **(*E-H)*** and anti-b-actin antibodies (loading control). For (*A-D*) data is presented as Hsp27 normalized levels to vehicle-treated cells. Panel (*A*) is repeated from main Fig. 1*C*. **** *p* < 0.0001, one-way ANOVA, followed by Tukey’s multiple comparisons test.

**Figure 4.**
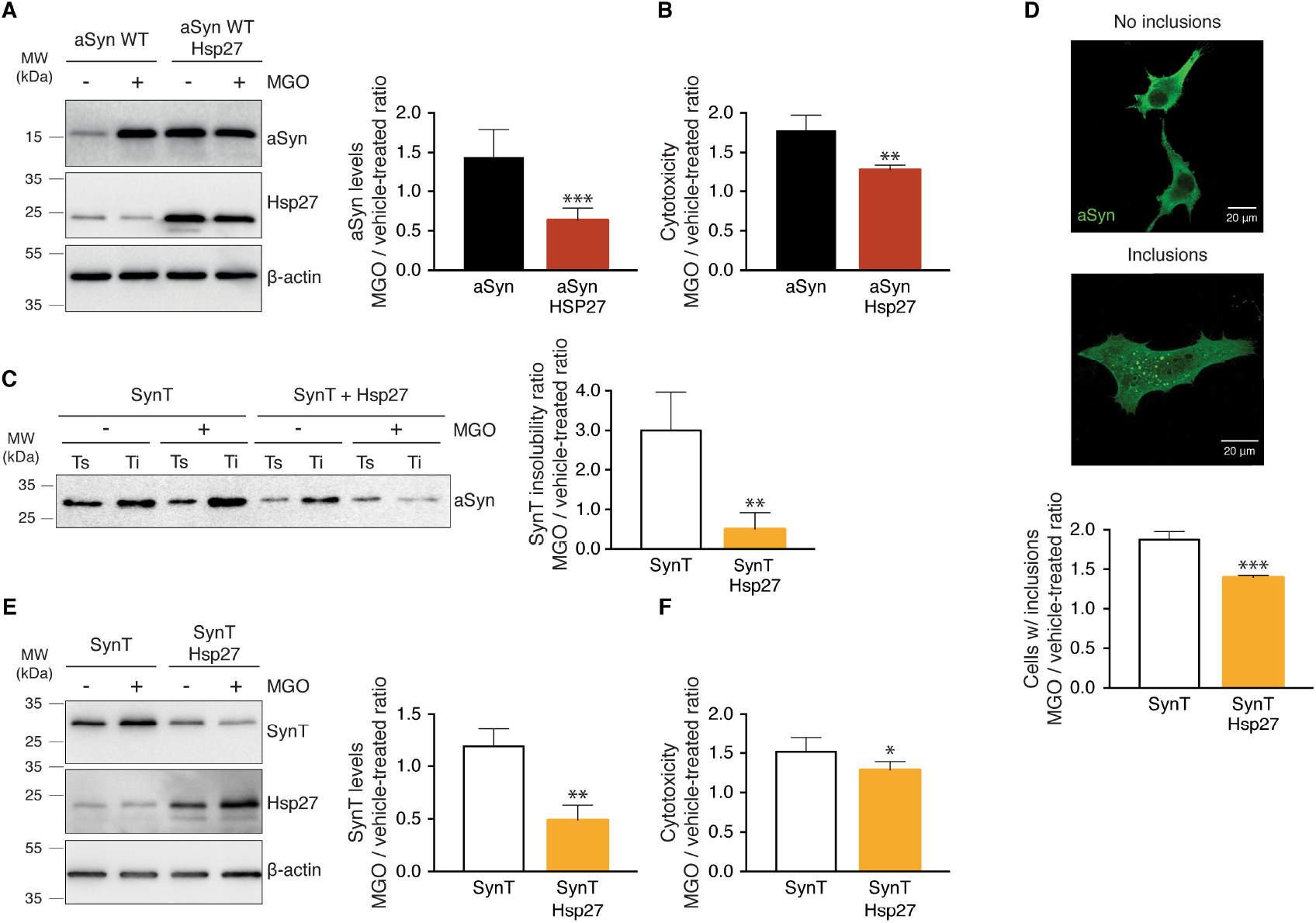
Hsp27 overexpression reduces cytotoxicity, insolubility and aggregation of aSyn in glycating conditions. Cells expressing aSyn or SynT, or co-expressing aSyn or SynT together with Hsp27 were treated 2 times for 2 consecutive periods of 24h with vehicle or MGO (0.2mM). **(*A*)** Protein levels of cells expressing aSyn were determined by immunoblotting with anti-aSyn, -Hsp27 or -b-actin antibodies. **(*B*)** Cytotoxicity was determined by loss of membrane integrity (LDH release). **(*C*)** Triton^™^ X-100 soluble and insoluble (TS and TI) fractions were probed for a-synuclein. The ratio between soluble and insoluble fractions is normalized to vehicle-treated ratio and presented as SynT insolubility (n = 4). **(*D*)** Cells were processed for immunocytochemistry (aSyn, green), and the percentage of cells with inclusions determined. **(*E*)** Protein levels and **(*F*)** cytotoxicity of cells expressing SynT were determined as in (*A*) and (*B*). Data presented as fold ratio between MGO and vehicle-treated cells, at least n=3. * *p* < 0.05, ** *p* < 0.01, *** *p* < 0.001 unpaired t-test with equal SD.

In order to assess whether Hsp27 modulated the MGO-induced aggregation of aSyn, we used an established cell model using an aggregation-prone variant of aSyn (known as SynT) ^19,64,66–68^. This model consists of a modified form of aSyn where the C-terminus is fused to a truncated, non-fluorescent fragment of GFP (known as SynT) ^36^, ^69^. Using this model, we have previously shown that MGO-treatment of cells expressing SynT increases the percentage of cells with aSyn inclusions (51% to 85%) ^19^. The aggregation status of SynT was assessed using Triton^™^ X-100 solubility assays ^19^. Briefly, H4 cells overexpressing i) SynT; or ii) SynT together with Hsp27, were treated with MGO as previously described. Protein extraction was performed in non-denaturing conditions, and protein extracts solubilized in 1% of Triton^™^ X-100. Soluble and insoluble fractions were separated by centrifugation and analyzed by western blot. We observed that MGO treatment induced an increase of SynT insolubility (Fig. 4 *C*). Remarkably, Hsp27 expression not only prevented SynT insolubility, but also potentiated its solubility in comparison to control conditions (**Fig. 4 *C***). SynT aggregation was also assessed by immunocytochemistry, as previously described ^19,66^. Using the same paradigm, H4 cells were stained for aSyn and the percentage of cells with aggregates determined. Consistently, Hsp27 expression prevented the MGO-induced SynT aggregation (~ 25%) (**Fig. 4 *D***). In cells overexpressing SynT, we observed that Hsp27 was also able to prevent the MGO-associated increase of SynT levels (~ 59%) (**Fig. 4 *E***) and cytotoxicity (~ 16%) (**Fig. 4 *F***).

## Discussion

Recent reports suggest that type II diabetes is an important risk factor for PD. In particular, protein glycation seems to play an important role in synucleinopathies ^16–18^. We previously showed that aSyn is a target of glycation, which potentiates its oligomerization, impairs its clearance and membrane binding ability, resulting in its accumulation, aggregation and cytotoxicity, inducing dopaminergic neuronal loss and motor impairments in animal models ^19^.

However, glycation can affect many other proteins in addition to aSyn, thereby contributing to a variety of cellular pathologies. To investigate this, we evaluated the pattern of glycation in a cellular model of synucleinopathies treated with increasing concentrations of MGO, as a model glycating agent since it is the most significant and highly reactive glycating agent in cells, mainly generated as a by-product of glycolysis ^70^. Surprisingly, although we were expecting an overall increase in the levels of glycated proteins, we observed dominant signal corresponding to a ~25 kDa protein whose levels decreased with increased levels of MGO. Using peptide mass fingerprinting we identified this protein as Hsp27 and confirmed this using a specific antibody. In fact, we found that the total levels of Hsp27 decreased in an MGO-dependent manner.

Hsp27 was previously shown to be protective in models of synucleinopathies. In particular, it reduces aSyn oligomerization ^23,49,71^ and prevents the toxicity of exogenous aSyn species in neuronal cells ^23,50^. Hsp27 reduces the aggregation of aSyn ^35^ and of other proteins associated with other neurodegenerative disorders ^33,72–74^. Therefore, we hypothesized that under glycating conditions, restoring or inducing the levels of Hsp27 might prevent the deleterious effects of cellular glycation in aSyn patohogenesis.

To test our hypothesis, we first compared the kinetics of recombinant aSyn oligomerization alone or in the presence of Hsp27, in control or glycating conditions. In contrast to other studies that tested aSyn:Hsp27 ratios of 1:1 or 5:1 ^23,49^, we performed our analysis in sub-stochiometric concentrations of 300:1 or 300:7, to assess the effect of much lower amounts of Hsp27, We confirmed our previous findings that glycation potentiates aSyn oligomerization. Moreover, we also validated that Hsp27 is able to reduce non-glycated aSyn aggregation. Remarkably, in the presence of Hsp27, the deleterious effects of glycation over aSyn oligomerization were almost completely abolished, avoiding the formation of stable HMW species. Also, although glycated aSyn species are cytotoxic, the species formed in the presence of Hsp27 were not. These findings provide important evidence that Hsp27 may directly suppress the pathogenicity of aSyn-glycation.

In order to further validate our hypothesis, we evaluated if Hsp27 overexpression would be protective in cell models of aSyn toxicity and aggregation. Impressively, Hsp27 alleviated the MGO-induced pathogenic phenotypes, reducing cytotoxicity in cells expressing WT aSyn or an aggregation-prone version of aSyn (SynT). Hsp27 significantly reduced the levels of aSyn or SynT and reduced the aggregation of aSyn.

Notably, the cytotoxic protection provided by Hsp27 in glycation conditions (28%) is higher than in non-glycation conditions (10%), suggesting a relevant role of Hsp27 in glycation conditions. Interestingly, MGO was shown to modify the α-crystallin domain of Hsps (namely, Hsp27) probably by exposing hydrophobic sites that would, otherwise, not be available for chaperone function. In addition, MGO might induce the formation of large oligomers of Hsp27, enhancing its activity ^51,75^. This enhanced Hsp27 activity might be responsible for reducing aSyn cytotoxicity, by reducing the formation of aSyn oligomers, or by preventing apoptosis, since Hsp27 glycation potentiates its anti-apoptotic potential, as described in several types of cancers ^54,55,76^. Although in the present study we could not confirm the formation of large oligomeric species of Hsp27, we hypothesize that the strong reduction of Hsp27 protein levels would prevail over a possible gain-of-function of Hsp27 as a molecular chaperone. The presence of Hsp27 in Lewy bodies suggests it may play a relevant role in PD ^77,78^. Actually, a decrease in the activity of Hsps and of other protein quality control components is associated with ageing and neurodegeneration ^79,80^. Molecular chaperones are crucial for stabilising protein conformations and for directing misfolded or damaged proteins for degradation by the different clearance pathways.

Therefore, our work highlights the importance of Hsp27 in suppressing the pathological conditions induced by glycation (**Fig. 5**) and suggests that Hsp27 may constitute a suitable target for intervention in synucleinopathies and other neurodegenerative disorders.

**Figure 5.**
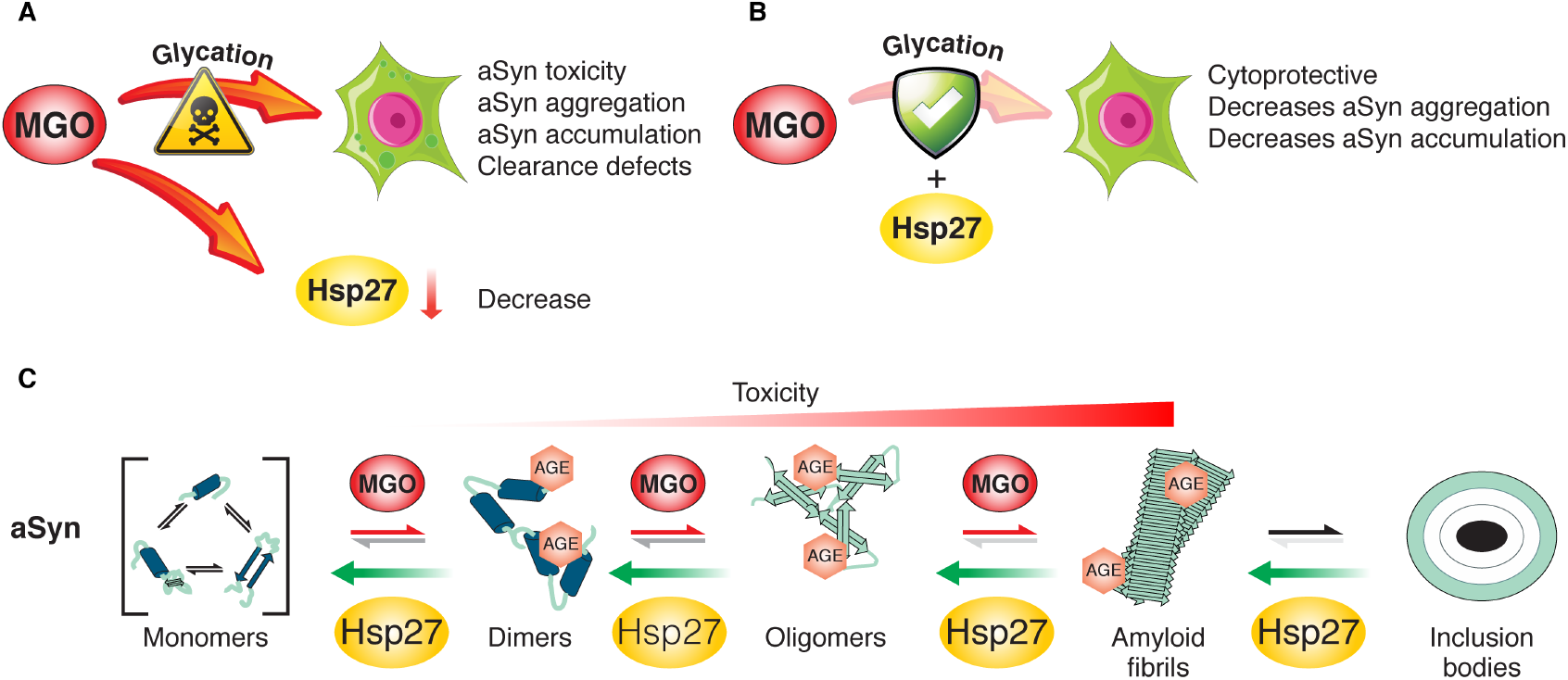
Model for the effects of Hsp27 on aSyn aggregation. **(*A*)** MGO-induced glycation exacerbates aSyn pathogenicity. Here, we showed that the levels of Hsp27 decrease in an MGO dependent manner. **(*B*)** Restoring Hsp27 expression prevents the deleterious effects of glycation in a cellular model of synucleinopathies, reducing aggregation both in vitro and in cells. **(*C*)** Based on our aSyn aggregation studies, we hypothesize that Hsp27 might reduce the oligomerization of aSyn and, therefore, its toxicity.

## Acknowledgments

This study was supported by Fundação para a Ciência e Tecnologia (FCT) PTDC/NEU-OSD/5644/2014 and EXPL/NEU-OSD/0606/2012. Authors were supported by: HVM (FCT, SFRH/BPD/109347/2015); MO (FCT, EXPL/NEU-OSD/0606/2012); AC (FCT, PD/BD/136863/2018; ProRegeM – PhD programme, mechanisms of disease and regenerative medicine); BFG (PTDC/NEU-OSD/5644/2014). TFO is supported an EU Joint Programme - Neurodegenerative Disease Research (JPND) project (aSynProtec). The project is supported through the following funding organisations under the aegis of JPND - http://www.jpnd.edu (BMBF).

## Author Contributions

H. Vicente Miranda and T.F. Outeiro designed research; H. Vicente Miranda, A. Chegão, M. Oliveira and B.F. Gomes performed research; F.J. Enguita contributed new reagents or analytic tools; H. Vicente Miranda, A. Chegão, M. Oliveira and T.F. Outeiro analyzed data; H. Vicente Miranda, A. Chegão and T.F. Outeiro wrote the manuscript.

