## Supplemental table 1 for "Hsp27 reduces glycation-induced toxicity and aggregation of α-synuclein"

### TITLE

**Supplemental Table 1**

**Hsp27 identification by Peptide mass fingerprint (PMF) (trypsin digestion).** Theoretical peptide mass (Da); measured M/z (Da); deviation (ppm); Start-end Identified peptides; peptide sequences; putative glycated residues. Oxid (M); N-terminal acetylation; carbamidomethyl (M); carboxyethyl - CEL (K); argpyrimidine (R); hydroimidazolones - MG-H (R); tetrahydropyrimidine - THP (R); as variable modifications.

| Theoretical peptide mass (Da) | Measured M/z (Da) | Deviation (ppm) | Peptide Sequence | Peptide sequence and modifications | Glycated Residues |
|---|---|---|---|---|---|
| 1638.8785 | 1638.877 | -0.9 | [1-12] | MTE**RR**VPFSLLR [1xArgpyrimidine + 1xMG-H] | R4 + R5 |
| 987.6098 | 987.609 | -0.8 | [5-12] | RVPFSLLR | |
| 831.5087 | 831.509 | 0.4 | [6-10] | VPFSLLR | |
| 961.4526 | 961.453 | 0.4 | [13-20] | GPSWDPFR | |
| 1982.8933 | 1982.884 | -4.7 | [13-27] | GPSWDPF**R**DWYPHSR [1xArgpyrimidine] | R20 |
| 1163.6208 | 1163.622 | -0.7 | [28-37] | LFDQAFGLPR | |
| 2206.0636 | 2206.091 | 12.4 | [80-96] | QLSSGVSEI**R**HTAD**R**WR [1xMG-H + 1xTHP] | R89 + R94 |
| 2206.1291 | 2206.091 | -17.3 | [95-112] | W**R**VSLDVNHFAPDELTVK [1xArgpyrimidine] | R96 |
| 1146.6365 | 1146.631 | -4.8 | [113-123] | TKDGVVEITGK | |
| 1218.6576 | 1218.65 | -6.2 | [113-123] | T**K**DGVVEITGK [1xCEL] | K114 |
| 1793.8388 | 1793.808 | -17.2 | [128-141] | QDEHGYIS**R**CFTRK [1xMG-H] | R136 or R140 |
| 1796.8497 | 1796.846 | -2 | [128-141] | QDEHGYISRCFTRK [1xCarbamidomethyl] | |
| 1905.9916 | 1905.989 | -1.4 | [172-188] | LATQSNEITIPVTFESR | |
